# DataJoint Elements: Data Workflows for Neurophysiology

**DOI:** 10.1101/2021.03.30.437358

**Authors:** Dimitri Yatsenko, Thinh Nguyen, Shan Shen, Kabilar Gunalan, Christopher A. Turner, Raphael Guzman, Maho Sasaki, Daniel Sitonic, Jacob Reimer, Edgar Y. Walker, Andreas S. Tolias

## Abstract

A new resource—*DataJoint Elements*—provides modular designs for assembling complete workflow solutions to organize data and computations for common neurophysiology experiments. The designs are derived from working solutions developed in leading research groups using the open-source DataJoint framework to integrate data collection and analysis in collaborative workflows.

## Meeting challenges of advanced neurotechnologies in team science

We announce *DataJoint Elements*: a resource for open-source software modules for organizing data and computations for the major types of neurophysiology experiments. The project aims to provide an efficient approach to managing scientific data workflows—the multi-step methods for data collection, preparation, processing, and analysis—that researchers must perform in the course of an experimental study.

For defining and deploying the workflows, the project relies on DataJoint: the open-source library for MATLAB and Python enabling streamlined interactions with relational databases and bulk data stores [1]. By formalizing computational dependencies as a fundamental concept of its data model, DataJoint serves as an effective scientific workflow management system [2]. Similar to other formal scientific workflow management systems widely used in bioinformatics, astronomy, and other data-intensive disciplines, DataJoint automates computations in sequential activities for data collection and analysis [3–6].

For the past several years, a number of collaborative neuroscience groups have used DataJoint to define and automate their shared data workflows. Among them are IARPA’s MICrONS, Mesoscale Activity Project, the International Brain Lab, Princeton U19 (“Brain CoGS”), Columbia U19 (Team “MoC3”), Harvard U19 (Team “Dope”), Moser Group, and others. These consortia conduct high-quality research by pooling resources and expertise, establishing common management practices, adopting efficient data sharing standards, and developing shared computing infrastructure [7]. As many as a hundred individual labs already use DataJoint in diverse data modalities such as multielectrode electrophysiology, calcium imaging, light and electron microscopy, behavior tracking, sensory stimulation, transcriptomics, and optogenetics. In these complex data ecosystems, DataJoint enables groups of scientists to collaborate with clear workflows while sharing data continuously and maintaining data integrity.

Many of these teams publish their DataJoint pipelines under open-source licenses so that working solutions can be found for most data modalities and experiment paradigms in neurophysiology. Yet new DataJoint users may struggle to find good starting points and didactic examples because the published solutions are often obfuscated with project-specific complexity while standardized, easily adopted end-to-end solutions are yet to emerge.

DataJoint Elements are derived from developments in leading neuroscience projects, integrating data acquisition and analysis tools developed by numerous academic projects. Essential motifs are extracted from projects across various neurophysiology modalities into a collection of simple modules ready for assembly into complete workflows for new projects in new combinations.

Experiment workflows are products of scientific research in their own right, essential to the reproducibility of scientific findings. The sharing and dissemination of scientific workflows is a key aspect of open science and FAIR principles, making output of scientific research Findable, Accessible, Interoperable, and Reproducible [8,9]. From this perspective, DataJoint Elements is the first systematic effort to compile and disseminate complete workflows for neurophysiology experiments, integrating essential neuroinformatics tools, resources, and interfaces, deployable on diverse IT infrastructure. The project allows researchers to adopt, customize, and extend complete workflows from existing elements already validated in previous research and to contribute their new developments back to the resource.

## More than FAIR: data integrity, flexible queries, and distributed computing

### Principles of effective data management in shared workflows

DataJoint is a library in Python and MATLAB for interacting with relational databases. Its key innovation is the integration of computational dependencies into its data model, producing an effective workflow management system.

Using DataJoint, scientists define and operate custom data pipelines for data collection and analysis directly from the same programming environments used for data analysis (Fig. 1). The diversity and complexity of the underlying data infrastructure is hidden from the scientist who only interacts with a uniform representation of the data pipeline regardless of the underlying data infrastructure. The data pipeline is visualized as a graph of nodes where each node represents a table of data and a class in MATLAB or Python providing methods for data queries and for automated computations within the pipeline. Thanks to a built-in data serialization framework, each table can accommodate complex data across modalities such as images, movies, electrophysiological traces. A typical infrastructure underlying DataJoint workflows includes a relational database system in combination with a bulk storage system, *e.g*. network-attached file storage or object storage such as Amazon’s Simple Storage System (S3).

**Figure 1:**
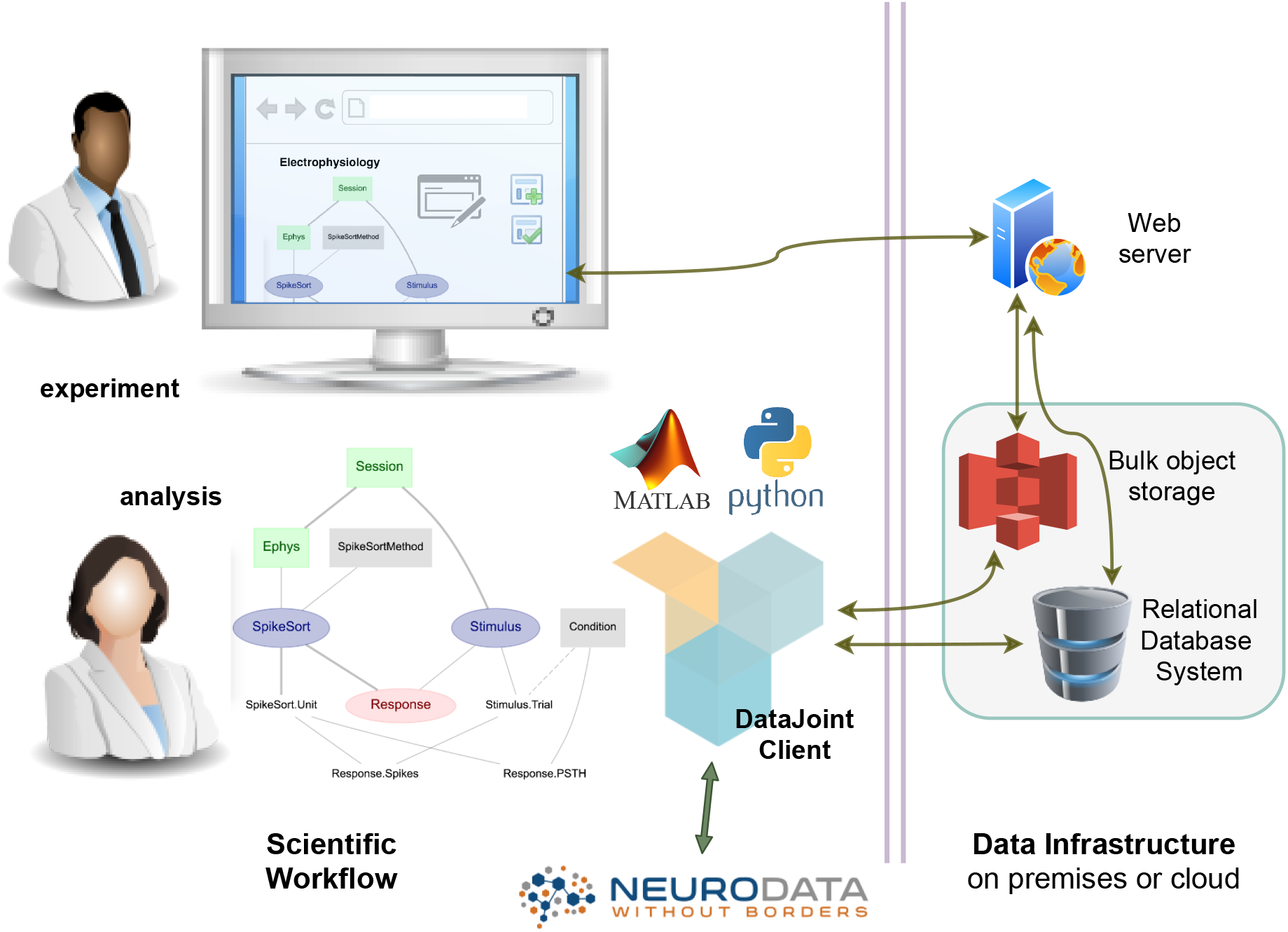
Scientists use DataJoint from their programming environments (MATLAB or Python) to define the structure of the data pipeline and its computational workflow. The data infrastructure (separated on the right) may be managed on premises or in the cloud. Web interfaces offer convenient interaction with the data pipeline. Diverse configurations are supported.

Computational workflows are modeled as directed acyclic graphs where each node is a table in a relational database containing or tracking stored data (Fig. 2A). Each node is linked with a class in a scientific programming language such as MATLAB or Python which builds the database table contents by supplying the necessary upstream data to user-defined computation code (Fig. 2B).

**Figure 2:**
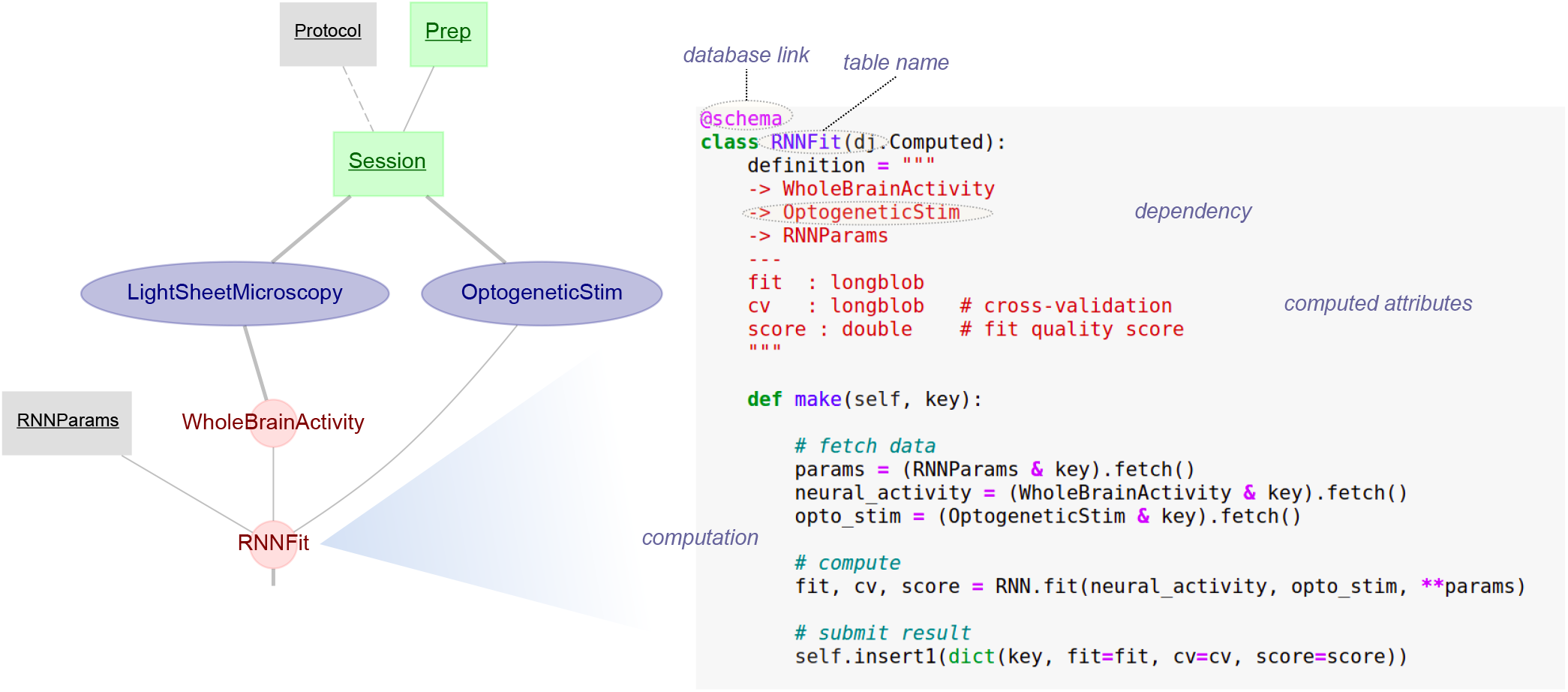
DataJoint serves as a framework for defining and executing scientific data workflows. It combines principled data management, workflow execution, and flexible queries accessed directly from the popular scientific programming languages MATLAB and Python. A) The graph represents the workflow for a neuroscience experiment where each node is a workflow step with a corresponding table in the database to store its data. Each node also has a class in the host programming language for scientific computing that defines the node and any computations that must be performed to populate the table. B) The Python class definition for one of the nodes in the workflows, RNNFit, defines the callback function, make, which defines the computation.

DataJoint solves several key technical challenges for using databases effectively in scientific workflows, in particular:

**Complete relational data model** DataJoint allows programming databases directly from a scientific computing language such as MATLAB and Python without the need for programming in SQL.
**Data definition language** DataJoint’s simple data definition language allows creating database schemas with tables and dependencies between them directly from the scientific language.
**Diagramming notation** DataJoint allows visualizing and navigating the database structure graphically directly from MATLAB and Python.
**Query language** DataJoint’s data query language creates flexible and precise queries using only a few operators.
**Serialization framework** DataJoint allows storing and retrieving numerical arrays and other scientific data structures directly in the database in a language-independent way, enabling the use of different programming languages on the same data pipeline.
**Automated distributed computations** DataJoint allows defining computations as part of its data model and provides a built-in job reservation process for computations on distributed compute nodes.

The FAIR principles govern the sharing of static datasets at the outcome of a project (Fig. 3). Thanks to its principled data model, integrated computational dependencies, and the use of relational database systems to structure the data and to orchestrate computations, DataJoint supports additional principles for data management suitable for sharing live data and computations in the active phase of a dynamic collaborative project:

**Data structure** DataJoint defines, communicates, and enforces the structure of the workflow in the form of a relational schema, ensuring that all required data are complete and in proper form before allowing the next step in the workflow. This structure reflects the logic of the experiment study, evolving with the project.
**Data integrity** DataJoint helps to ensure unique mapping between real-world entities and their digital representations (entity integrity), correct associations among entities (referential integrity), and inseparability of parts making up a single entity (compositional integrity). These guarantees prevent unintended data corruption, incompleteness, loss, misidentification, or mismatch.
**Data consistency** When multiple agents, both human and digital, access the same data concurrently, for both reading and writing, then integrating their activities with minimum delay and without violating data integrity becomes challenging. Data consistency, supported through the use of serializable transactions and consensus algorithms in modern distributed database systems, guarantees that digital representations of real-world entities appear the same to all agents with all changes correctly applied. DataJoint makes use of these features to achieve data consistency under the pressures of concurrent access by multiple agents in dynamic team projects.
**Workflow** DataJoint schemas are structured and visualized as directed acyclic graphs (DAGs), simultaneously representing the database schema, the data pipeline, and the workflow as a sequence of operations in conducting a study.
**Performance** DataJoint databases allow creating indexes to accelerate common queries, avoiding traversal of entire datasets when finding entities with specific attributes.
**Precise queries** Data is rarely consumed in the same form as it is deposited. Database queries allow transforming the data into a desired form remotely, without the need for transferring entire datasets for specific analysis.
**Distributed computing** A workflow management system must support a mechanism for orchestrating and automating computations by multiple nodes. DataJoint provides a built-in process for querying and reserving available jobs. Automation of computations leads to increased productivity in research teams.

**Figure 3:**
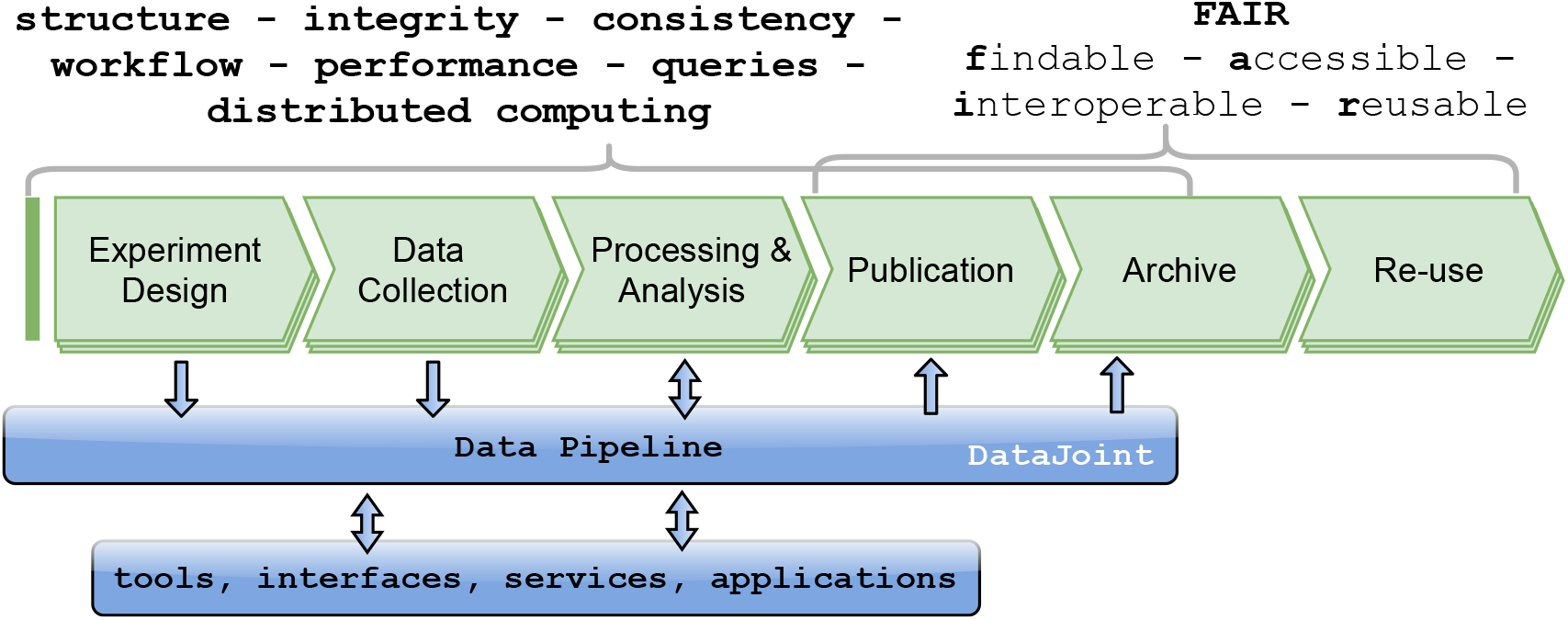
Data management considerations across the lifecycle of projects featuring shared data pipelines.

### Good workflow management fosters open science

Open science for integrity and reproducibility of research is a major thrust of all major biomedical research initiatives and the BRAIN Initiative in particular with data sharing recognized as a critical need for the effective use of research funds and resources [10]. However, today scientists incur a substantial cost in making their science open due to the heavy burden in preparing code and data for sharing or even in reusing resources shared by others [11, 12]. The lack of structure and organization of data management within the project itself makes it difficult to compile all the data to comply with FAIR standards. A student rushing to defend her thesis or a postdoc applying for academic positions can ill afford the effort of transforming their data and code into a form suitable for sharing. With DataJoint, all the data is already complete, well-structured, and ready for export during the active phase of the project.

### Going big: roles and responsibilities in a data-centric collaboration

DataJoint workflows often start small, running on a graduate student’s laptop. But they can also quickly evolve and scale to entire labs, and then to multi-lab consortia.

In planning data-rich, computation-intensive projects, we distinguish four layers of technical challenges and areas of expertise: *Experiment Workflow, Data Science, Data Engineering*, and *IT Infrastructure* (Figure 4). Each of these areas requires a principled and integrated approach, allowing for separation of roles and responsibilities. At the top level, experimentalists are mainly concerned with efficiently defining and executing their *experiment workflows*—the end-to-end sequences of steps necessary for completing a study: tracking all elements of the experiment, entering and acquiring data, synchronizing different modalities, processing and analysis, and generating visualizations, statistical tests and reports, publishing, sharing, and archiving.

**Figure 4:**
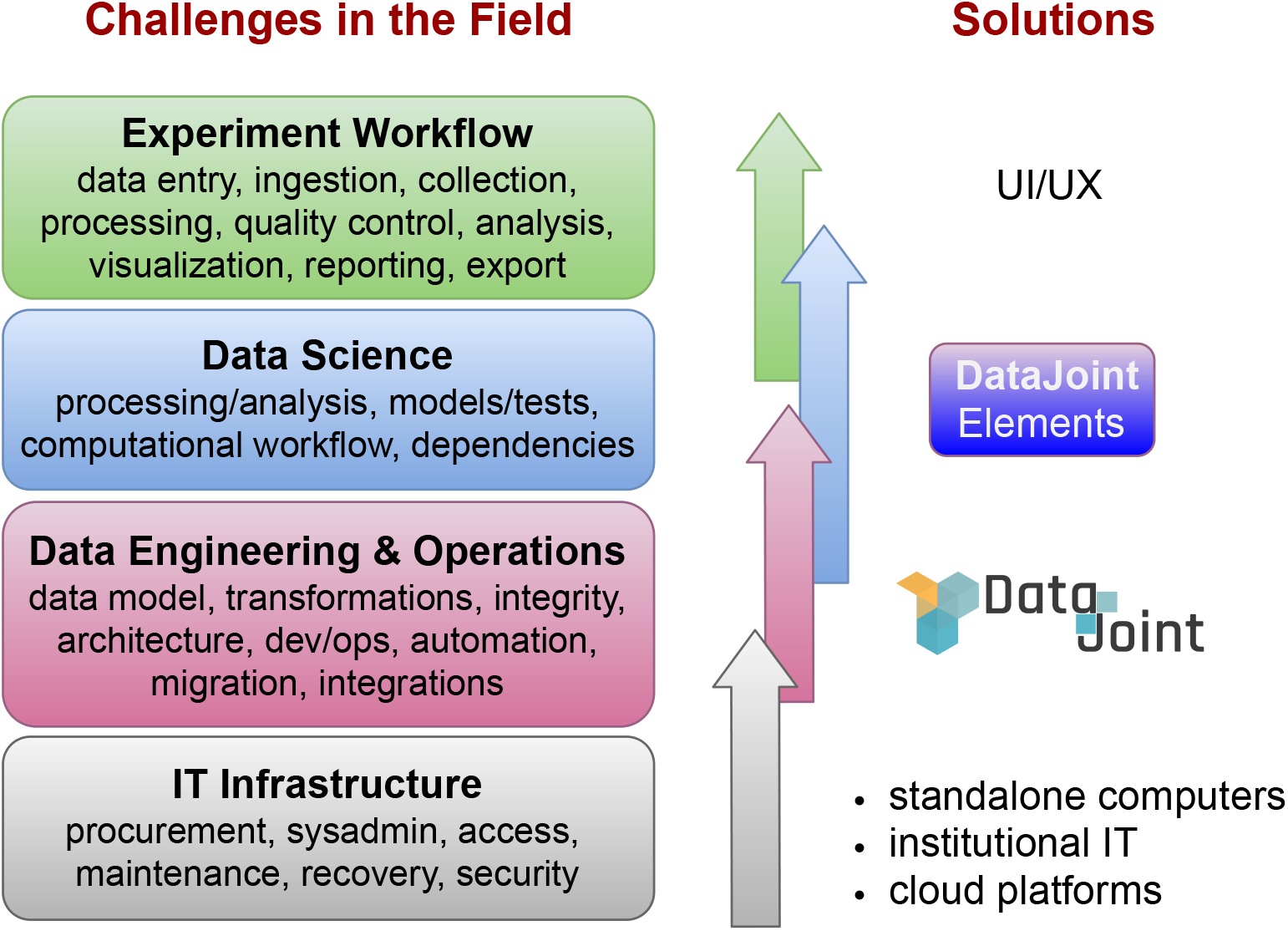
Challenges in planning data-intensive projects and their solutions in the DataJoint ecosystem.

A level down from experiment workflows are the *Data Science* tools for processing and analyzing the data. Most of these tools are developed as academic research projects or as custom solutions in individual labs. These tools are commonly shared across the community in the form of stand-alone open-source applications or code. Then every lab still needs to solve a similar set of *Data Engineering and Operations* problems to integrate these tools into the particular workflows of their experiments: user interfaces for data entry, scripts for data acquisition and processing, sequences of computing steps, data management processes, and visualization programs, addressing the challenges of data integrity in data transformations. Finally, these activities must be mapped onto the available *IT Infrastructure*, which can range from an individual laptop, to lab computer networks, institutional data centers, and public cloud platforms.

Clear separation of roles under a common frame-work enables increased the productivity and efficiency of research activities in larger projects.

## Progress and Development Plan

### Project Management

DataJoint Elements is led by the DataJoint NEURO team listed among the co-authors of this paper with Dimitri Yatsenko as principal investigator, with funding and programmatic support provided by the National Institute Of Neurological Disorders And Stroke of the National Institutes of Health. A Scientific Steering Group comprising systems and computational neuroscientists with diverse backgrounds provides project oversight and guidance (See Acknowledgements).

### Resource components

The Resource developed under this project aims to provide an efficient approach for neuroscience labs to create and manage scientific data workflows for the major modalities of neurophysiology experiments using advanced instrumentation and state-of-the-art analysis. The Resource comprises two major components:

**The DataJoint framework:** the open-source framework for data pipelines and automated computational workflows, including related documentation and tutorials, interfaces, deployment tools, and utilities. The central website for DataJoint is https://datajoint.io.
**DataJoint Elements:** a collection of curated modules for assembling workflows for the major modalities of neurophysiology experiments. These modules are designed for modularity to allow flexible use in various combinations. They integrate automated ingestion of acquired data, common data processing and analysis tools, neuroinformatics resources, and provide related utilities. The central website for the Resource is https://elements.datajoint.io, listing the project’s policies, licensing, quality assurance, outreach, and dissemination plans. It provides references for all released components.

### Project selection

We have adopted a set of criteria for accepting and prioritizing new components for development and support in the Resource. Development for a new experimental modality begins by engaging with a research group who have already developed a functioning DataJoint workflow as an open-source project. Our team reviews the workflow design and interviews the team about their process for conducting experiments and analyzing data. For all third-party tools or resources included in the proposed component, their long-term maintenance roadmap must be established. When possible, we will contact the developer team to establish a joint sustainability roadmap or to identify alternative solutions as replacement. The project is accepted for development if significant interest in its experiment modality can be established, with 100+ labs projected to use the modality over the next few years.

### Validation and Quality Assurance

After the initial development and internal testing, all new components will be first released in Alpha. During this phase, we engage external research teams to test and validate the complete workflows in real-life experiments with our team’s engineering support. During this phase, significant design changes may be performed and not all features may be completely developed. However, several features should be usable and suitable for testing and validation.

After the initial validation phase, the workflows are made available to the general public with a warning of Beta status and that the released code may be subject to errors and changes. During this phase, feature developments are complete with a focus on collecting user feedback to make design improvements and bug fixes. After collecting feedback and fixing any issues, our team will provide support to address any discovered issues and upgrade the workflows for new instrumentation and analysis tools, following the same quality assurance process as the initial release.

### Current developments

The first batch of DataJoint Elements to be released by the Resource provide experiment workflows for array electrophysiology, multi photon calcium imaging, and Miniscope calcium imaging. These workflows are currently in Alpha release undergoing validation in several neuroscience labs. The Beta release is planned for May 1, 2021, along with documentation and training materials.

#### Lab, Animal, Genotyping, and Session

Most workflows begin with information about the lab describing experiment setups, animal subjects, and experiment sessions. The Lab Element, Animal Element (with an optional genotyping module), and Session Element provide typical implementations of these schemas. Their code repositories are:

- https://github.com/datajoint/element-lab
- https://github.com/datajoint/element-animal
- https://github.com/datajoint/element-session

#### Array Electrophysiology

The Element is designed to accommodate the growing number of labs using Neuropixels probes versions 1.0 and 2.0 [13, 14]. The workflow is designed to ingest data acquired by the OpenEphys [15] or SpikeGLX [16] acquisition software and to isolate spiking units using the Kilosort algorithm. The design is derived from DataJoint electrophysiology pipelines developed by the International Brain Lab, Mesoscale Activity Project (HHMI/Janelia and BCM).

The development timeline and code repository for this Element can be found at

- https://github.com/datajoint/element-array-ephys.

#### Calcium imaging

This Element is designed to process laser-scanning multiphoton imaging of calcium signals. The workflow is designed to ingest data acquired by ScanImage (Vidrio Technologies) and Scanbox acquisition software (https://scanbox.org). The workflow integrates Suite2p [17] and CaImAn [18] analysis tools to perform motion correction, segmentation, fluorescence trace extraction, spike inference, and cell classification. The design is derived from calcium imaging pipelines developed by Andreas S. Tolias’ Lab (BCM) and BrainCoGs labs (Princeton University).

Figure 5 depicts the schema diagram of the Calcium Imaging Element as part of a larger workflow, also showing processed experiment data contained within the tables.

**Figure 5:**
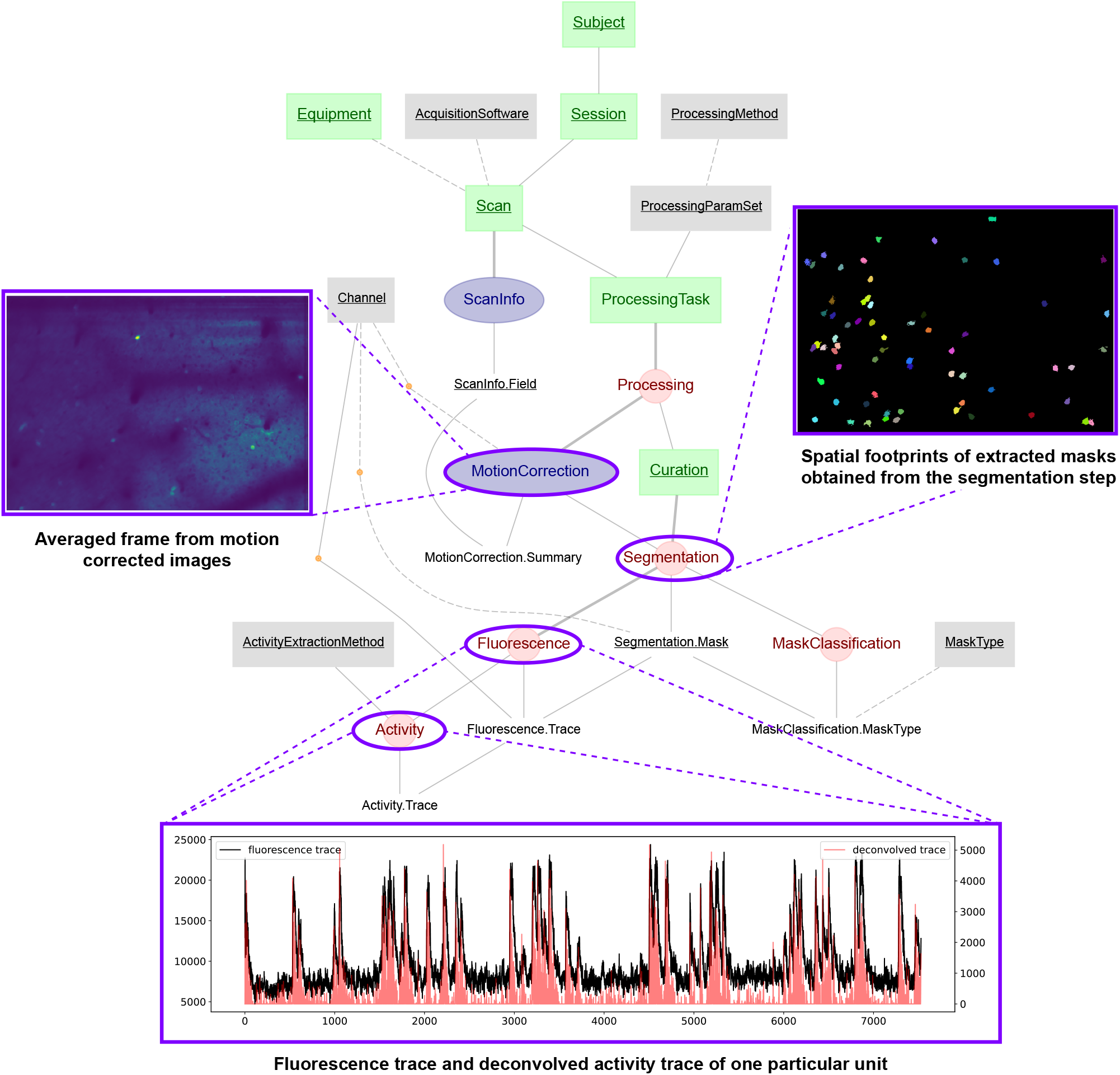
The Calcium Imaging Element embedded in a larger workflow. It performs data ingest, motion correction cell segmentation and classification, fluorescent trace extraction, and calcium spike inference (deconvolution), and support for multiple manual curations of the segmentation results.

The project information and code repository for this Element can be found at

- https://github.com/datajoint/element-calcium-imaging.

#### Miniscope Imaging

This Element is designed to accommodate the special acquisition and processing involved in calcium imaging using the UCLA Miniscope imaging probes, with future plans for supporting Inscopix probes as well. Due to its versatility, this modality is among the fastest growing, with about 700 labs using the UCLA variant alone. The workflow ingests data from Miniscope DAQ V3 acquisition software and several data analysis tools.

The project information and code repository for this Element is

- https://github.com/datajoint/element-miniscope.

### Utilities and training infrastructure

Along with the Elements, the Resource continues to develop the DataJoint framework itself, driven by the needs of specific projects. This includes general utilities, interfaces, and training resources.

This includes:

**Pharus:** a REST API for building custom applications to interact with DataJoint pipelines.
  - https://datajoint.github.io/pharus/

**DataJoint LabBook:** a frontend application for entering and querying data in data pipelines.
  - https://github.com/datajoint/datajoint-labbook/

**DataJoint Playground:** An online platform for interactive exercises to learn to work with DataJoint pipelines in a series of Jupyter-based tutorials. This infrastructure will be used to deliver upcoming workflow-specific training courses, including sample data.
  - https://playground.datajoint.io

### Future milestones and roadmap

With the first batch of products approaching their release in June, 2021, the project is preparing the milestones for Year 2.

These milestones include the workflows for common neurophysiology experiment modalities:

- Trial-based behavior
- Behavior tracking / pose estimation
- Trial-based stimulation: visual
- Trial-based stimulation: optogenetics

Additional Elements will track other experiment information:

- Animal procedures
- Synchronization between modalities
- Quality control: electrophysiology
- Quality control: calcium imaging

The following utilities are also considered:

- Data export and packaging for sharing and publishing
- Integration with data publishing platforms and archives

We are actively engaging precursor projects already using variations of these solutions, developers of the analysis tools, and validation sites.

The Resource has a five-year roadmap with a sustainability plan beyond the initial funding period, with annual assessments of adoption rates, trends. Besides covering the major modalities of neurophysiology experiments, the Resource will focus on integration with the broader neuroinformatics infrastructure, resources, and standards such as brain atlases, coordinate frameworks, ontologies, and nomenclatures.

## Acknowledgements

Research reported in this publication was supported by the National Institute of Neurological Disorders and Stroke of the National Institutes of Health under Award Number U24NS116470. The content is solely the responsibility of the authors and does not necessarily represent the official views of the National Institutes of Health.

We thank the project’s *Scientific Steering Group*: Drs. Nicholas Steinmetz, John Cunningham, Mackenzie Mathis, Carlos Brody, Karel Svoboda, and Loren Frank.

We thank the projects that provide their opensource workflows, providing the source material for DataJoint Elements:

- Andreas S. Tolias’ Lab (BCM, Project MICrONS)
- The International Brain Lab — https://internationalbrainlab.org
- Mesoscale Activity Project — Nuo Li (BCM) and Karel Svoboda (HHMI/Janelia)
- Princeton U19 — BRAIN CoGS
- Columbia U19
- Moser Group (Kavli)

We are grateful to the following groups and individuals who participated in the validation of the Elements in their experiments:

- Peyman Golshani’s Lab (UCLA)
- Anne Churchland’s Lab (UCLA)
- Andreas S. Tolias’ Lab (BCM)
- Efthymia (Mika) Diamanti, Manuel Schottdorf, Alvaro Luna, Adrian Bondy, Carlos Brody (Princeton)

We also thank the developers of the many state-of-the-art open-source analysis tools that we have integrated into the workflows in the Resource and for meeting with us to consult on the development roadmap:

- Daniel Aharoni (UCLA)
- Jinghao Lu (Fan Wang’s Lab, MIT)
- Biafra Ahanonu (UCSF)
- Carson Stringer and Marius Pachtariu (HHMI/Janelia)
- Andrea Giovannucci (University of N. Carolina)

## Notes

https://elements.datajoint.io

### Competing Interest Statement

The authors have declared no competing interest.

### Summary of Updates

Author affiliations updated; resource website updated.

## References

[1] D. Yatsenko et al. DataJoint: managing big scientific data using MATLAB or Python. bioRxiv, pp. 031658, November 2015.

[2] D. Yatsenko et al. Datajoint: A simpler relational data model. arXiv preprint arXiv:1807.11104, 2018.

[3] E. Deelman et al. Workflows and e-science: An overview of workflow system features and capabilities. Future generation computer systems, 25(5):528–540, 2009.

[4] M. Atkinson et al. Scientific workflows: Past, present and future, 2017.

[5] J. Liu et al. A survey of data-intensive scientific workflow management. Journal of Grid Computing, 13(4):457–493, 2015.

[6] J. Vivian et al. Toil enables reproducible, open source, big biomedical data analyses. Nature biotechnology, 35(4):314–316, 2017.

[7] N. Bonacchi et al. Data architecture for a large-scale neuroscience collaboration. BioRxiv, pp. 827–873, 2020.

[8] M. D. Wilkinson et al. The FAIR Guiding Principles for scientific data management and stewardship. Scientific Data, 3:160018, March 2016.

[9] C. Goble et al. Fair computational workflows. Data Intelligence, 2(1-2):108–121, 2020.

[10] NIH strategic plan for data science. https://datascience.nih.gov/strategicplan, 2018.

[11] R. A. Poldrack. The costs of reproducibility. Neuron, 101(1):11–14, 2019.

[12] C. Allen and D. M. Mehler. Open science challenges, benefits and tips in early career and beyond. PLoS biology, 17(5):e3000246, 2019.

[13] J. J. Jun et al. Fully integrated silicon probes for high-density recording of neural activity. Nature, 551(7679):232–236, 2017.

[14] N. A. Steinmetz et al. Challenges and opportunities for large-scale electrophysiology with neuropixels probes. Current opinion in neurobiology, 50:92–100, 2018.

[15] J. H. Siegle et al. Open ephys: an open-source, plugin-based platform for multichannel electrophysiology. Journal of neural engineering, 14(4):045003, 2017.

[16] J. J. Jun et al. Real-time spike sorting platform for high-density extracellular probes with ground-truth validation and drift correction. bioRxiv, pp. 101030, 2017.

[17] M. Pachitariu et al. Suite2p: beyond 10,000 neurons with standard two-photon microscopy. BioRxiv, 2017.

[18] A. Giovannucci et al. Caiman an open source tool for scalable calcium imaging data analysis. Elife, 8:e38173, 2019.

